# Ocular tropism of SARS-CoV-2 with retinal inflammation through neuronal invasion in animal models

**DOI:** 10.1101/2022.04.17.488607

**Authors:** Gi Uk Jeong, Hyung-Jun Kwon, Hyun Woo Moon, Gun Young Yoon, Hye Jin Shin, Ji Soo Chae, Seong-Jun Kim, In-Chul Lee, Dae-Gyun Ahn, Kyun-Do Kim, Suresh Mahalingam, Young-Chan Kwon

## Abstract

Although ocular manifestations are commonly reported in patients with coronavirus disease 2019 (COVID-19), there is currently no consensus on ocular tropism of severe acute respiratory syndrome coronavirus 2 (SARS-CoV-2). To investigate this, we infected K18-hACE2 mice with SARS-CoV-2 using various routes. We observed ocular manifestation and retinal inflammation with cytokine production in the eyes of intranasally (IN) infected mice. An intratracheal (IT) injection resulted in virus spread from the lungs to the brain and eyes via trigeminal and optic nerves. Ocular and neuronal invasion were confirmed by an intracerebral (IC) infection. Notably, eye-dropped (ED) virus did not infect the lungs and was undetectable with time. Using infectious SARS-CoV-2-mCherry clones, we demonstrated the ocular and neurotropic distribution of the virus *in vivo* by a fluorescence-imaging system. Evidence for the ocular tropic and neuroinvasive characteristics of SARS-CoV-2 was confirmed in wild-type Syrian hamsters. Our data provides further understanding of the viral transmission; SARS-CoV-2 clinical characteristics; and COVID-19 control procedures.

**Summary:** SARS-CoV-2 can spread from the respiratory tract to the brain and eyes via trigeminal and optic nerves in animal models. This ocular tropism of SARS-CoV-2 through neuronal invasion likely causes ocular manifestation and retinal inflammation.

**Graphical Abstract:** 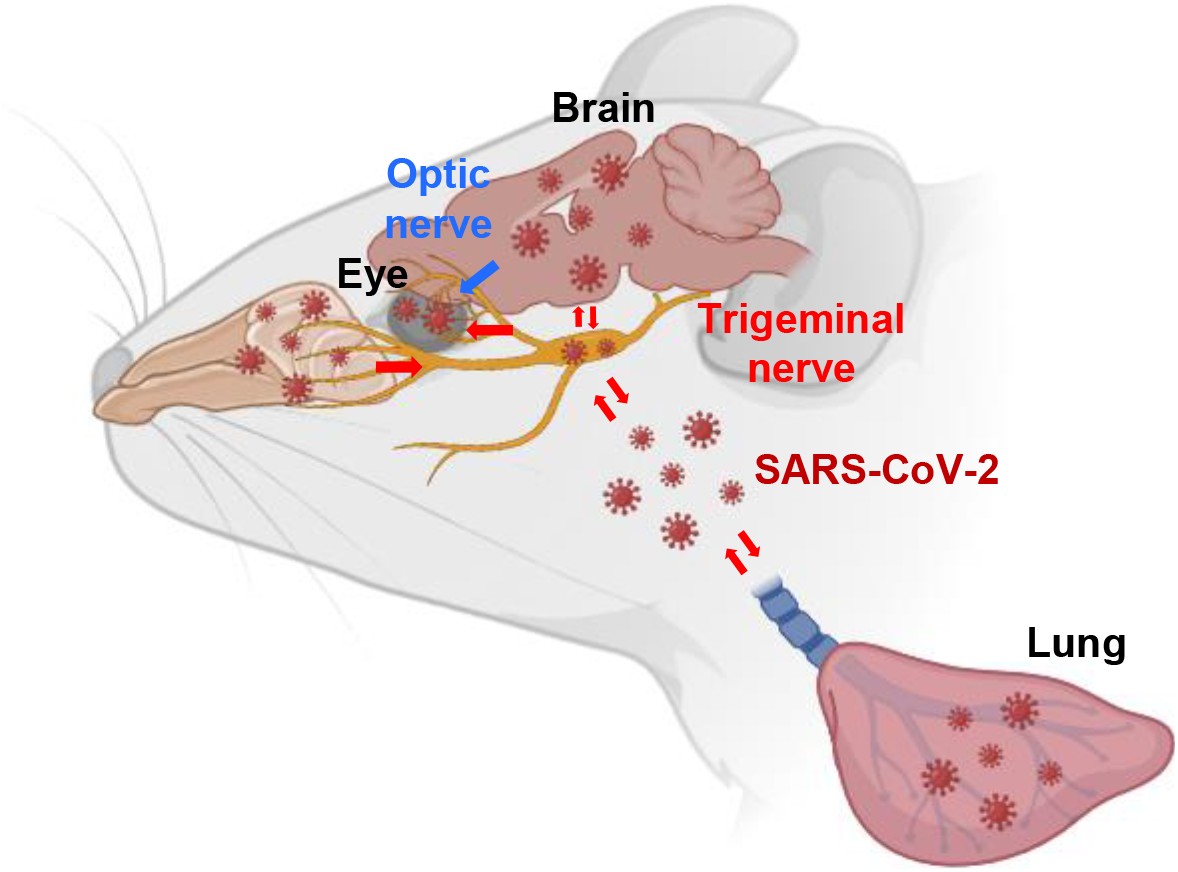

## Introduction

As of 9 January 2022, the ongoing the coronavirus disease 2019 (COVID-19) pandemic caused by severe acute respiratory syndrome coronavirus 2 (SARS-CoV-2) has threatened global public health with over 300 million confirmed cases and over 5 million deaths (https://www.who.int/emergencies/diseases/novel-coronavirus-2019/situation-reports). SARS-CoV-2 preferentially infects cells in the respiratory tract; however, it has a broad organotropism beyond the respiratory tract, affecting the kidney, small intestine, heart, and brain, and is believed to cause multi-organ dysfunction in patients with COVID-19 (Liu et al., 2021a; Puelles et al., 2020; Thakur et al., 2021).

Several respiratory viruses, such as species D adenoviruses and subtype H7 influenza viruses, possess an ocular tropism and cause ocular diseases in humans (Belser et al., 2013). While ocular manifestations and abnormalities have been commonly reported in patients with COVID-19 (Chen et al., 2020; Wan et al., 2021), the ocular tropism and pathologies of SARS-CoV-2 are still unclear. In some clinical and post-mortem cases, viral RNA was not detected by the PCR analysis of conjunctival swabs of three patients with COVID-19 with bilateral conjunctivitis (Pirraglia et al., 2020) and in 16 aqueous humor and vitreous samples (List et al., 2020). In contrast, a cross-sectional study in Italy reported that viral RNA was found on the conjunctival swabs of 52 of 91 patients with COVID-19, with a high rate of detection in severe patients (Azzolini et al., 2021). Other cross-sectional studies in China (Li et al., 2021) and Brazil (Gasparini et al., 2021) and post-mortem examinations (Penkava et al., 2021; Sawant et al., 2021) also reported the detection of viral RNA using conjunctival swabs or in the aqueous humor of the patients with or without ocular manifestations.

Given that the ocular surface represents an additional mucosal surface exposed to infectious aerosols, eyes are considered as a potential transmission route of SARS-CoV-2. Previous studies have supported this and have demonstrated SARS-CoV-2 receptor expression on the human ocular surface (Leonardi et al., 2020; Zhou et al., 2020a). Deng et al showed SARS-CoV-2 potential entry through the conjunctiva in rhesus macaques by conjunctival inoculation (Deng et al., 2020). In addition, SARS-CoV-2 can infect human embryonic stem cell-derived ocular epithelial cells (Eriksen et al., 2021) and human conjunctival epithelial cells (Singh et al., 2021). Thus, it needs to be elucidated whether eyes can be a primary or secondary virus entry site to devise preventative or therapeutic strategies against SARS-CoV-2 transmission.

In this study, we evaluated the ocular tropism and the possible ocular transmission of SARS-CoV-2 by using K18-hACE2 mice and wild-type Syrian hamsters. Through various inoculation routes, we demonstrated the ocular tropism of SARS-CoV-2 via neuronal invasion of trigeminal (TN) and optic nerves (ON) in the mouse model. Using infectious SARS-CoV-2-mCherry clones, we examined the ocular and neurotropic distribution of the virus *in vivo* by a fluorescence-imaging system. We provided evidence for the ocular tropic and neuroinvasive characteristics of SARS-CoV-2 in wild-type Syrian hamsters.

## Results

### Detection of SARS-CoV-2 and ocular manifestations in the eyes of infected mice

A previous study reported that Zika virus (ZIKV) infection of the eyes led to the development of conjunctivitis and panuveitis, and caused inflammation in the uveal tissues of a mouse model (Miner et al., 2016). To investigate whether the pathogenesis of the ocular disease is caused in response to the SARS-CoV-2 infection in K18-hACE2 mice, we intranasally (IN) challenged mice with approximately 10^4^ plaque-forming units (PFU) of the virus or the same volume of PBS (mock group). We observed about 20 percent of weight loss on the 7^th^ day post infection (dpi; Fig. 1A) and mortality on 8 dpi (Fig. 1B). Notably, on 6 dpi, tearing and eye discharge occurred in one quarter of the infected mice (Fig. 1C). We then evaluated the presence of SARS-CoV-2 in the eyes of the mice after the IN infection. On 6 dpi, the virus quantity in the lungs and eyes was assessed by plaque assay. The infectious viral titer of the eyes was as high as that of the lung (~10^6^ PFU/g; Fig. 1D). Given the detection of viral RNAs in tears derived from patients with SARS-CoV (Loon et al., 2004) and SARS-CoV-2 (Karimi et al., 2020), we assessed viral RNA copies in the lacrimal gland by RT-qPCR targeting the nucleocapsid gene on 6 dpi (Fig. 1E). Unexpectedly, the viral load of the lacrimal gland was significantly lower than that of the eyes and similar to that of the spleen (~10^4^ viral RNA copies per microgram total RNA), one of the tissues that was barely susceptible to the infection (Winkler et al., 2020). These results demonstrate the presence of the infectious viruses in the eyes, suggesting that the eyes are a target of SARS-CoV-2 infection.

**Figure 1.**
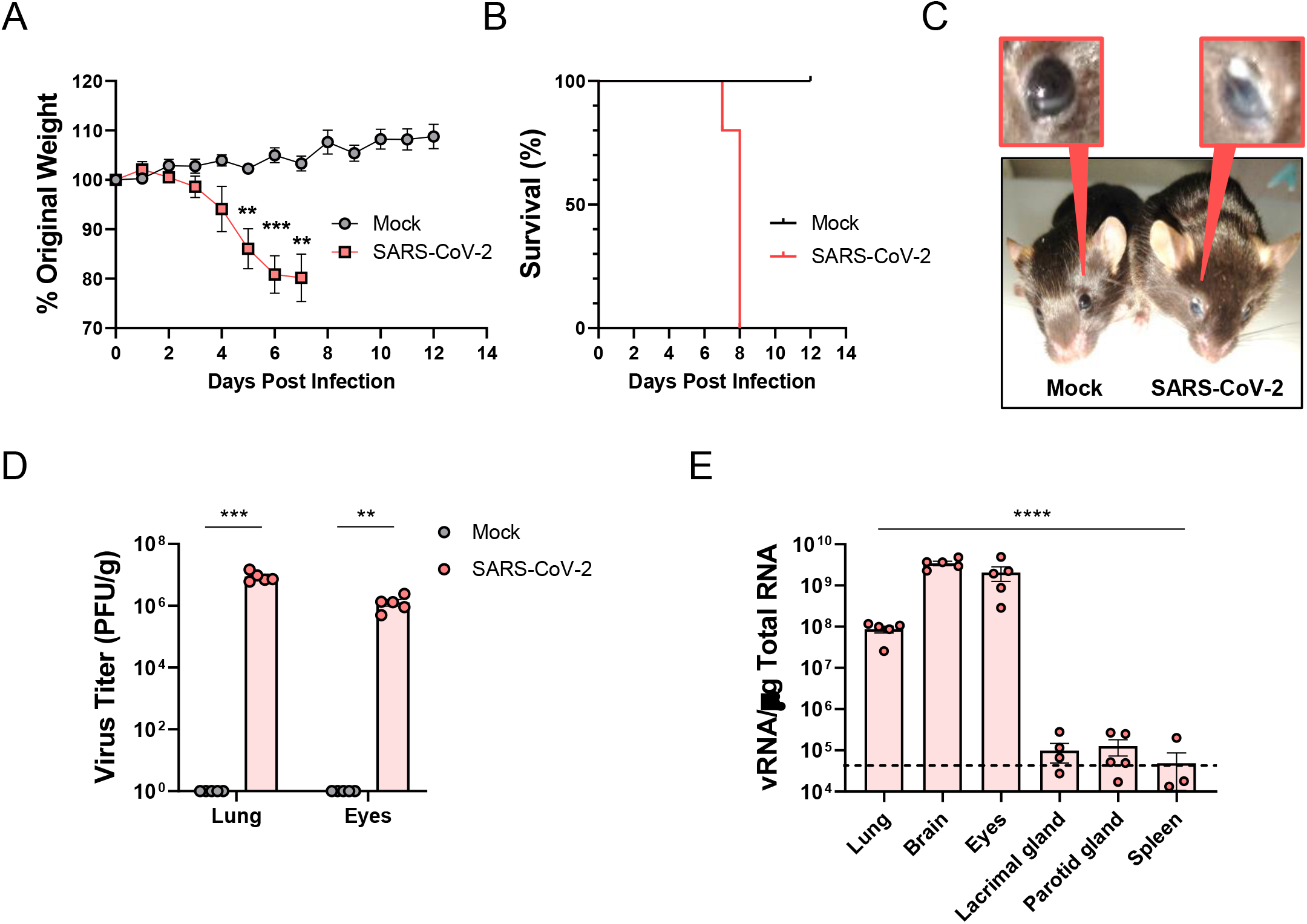
Clinical features and virus titers in the eyes of SARS-CoV-2-infected mice. Eight- to nine-week-old male K18-ACE2 mice were intranasally mock-infected or infected with approximately 10^4^ PFU of SARS-CoV-2 (*n* = 5 for mock-infected and infected mice, respectively). **(A)** Body weight changes shown as percentage of starting weight and **(B)** Survival was evaluated at the indicated dpi. **(C)** Representative image of tearing and eye discharge in SARS-CoV-2-infected mice (Right) compared with mock-infected mice (Left). **(D)** Viral load in the lungs and eyes was analysed by plaque assay. **(E)** Viral RNA levels in the lungs, brain, eyes, lacrimal gland, parotid gland, spleen, and colon were assessed by RT-qPCR. Viral RNA copies were cut-off by the limit of detection (10^4^ copies/μg). A dashed line indicates the viral RNA levels of spleen, barely susceptible to SARS-CoV-2 infection. Symbols represent means ± SEM. Statistically significant differences between the groups were determined by multiple t-test (A), Student’s t-test (D), or one-way ANOVA (E). \*\**P* < 0.01; \*\*\**P* < 0.001; \*\*\*\**P* < 0.0001. SARS-CoV-2: severe acute respiratory syndrome coronavirus 2; PFU: plaque-forming unit; dpi: days post-infection

### Retinal inflammation and cytokine production in the eyes of SARS-CoV-2-infected mice

We further examined histopathological and inflammatory changes in response to the infection using H&E-stained eye sections from mock- or SARS-CoV-2 IN infected mice. Since retinal thickness has been proposed and evaluated as a potential inflammatory marker (Steiner et al., 2019), we measured retinal thickness following the infection to estimate retinal inflammation. In comparison with mock group, apparent lesions in retinal tissues were observed in the SARS-CoV-2-infected mice (Fig. 2A). The mean of retinal thickness (from the ganglion cell layer to the nuclear layer) of mock mice was 46.27 μm and that of the infected mice was 75.14 μm (Fig. 2B). Viral infection significantly increased the retinal thickness by 1.62-fold and induced substantial accumulation of infiltrating inflammatory cells in ganglion cell and nuclear layers. Moreover, on 6 dpi, the multiplex immuno-analysis of the eyes showed elevated levels of pro-inflammatory chemokines and cytokines, including G-CSF, IP-10, MKC, MCP-1, MIP-2, and IL-12, in response to the infection (Fig. 2C). The levels of more pro-inflammatory chemokines and cytokines were significantly augmented in the brain, where the viral load was the higher than that of the eyes (Fig. 1E, Supplementary Fig. 1). These findings indicate that intranasally-administered SARS-CoV-2 promotes retinal inflammation and cytokine production in the eyes of K18-hACE2 mice.

**Figure 2.**
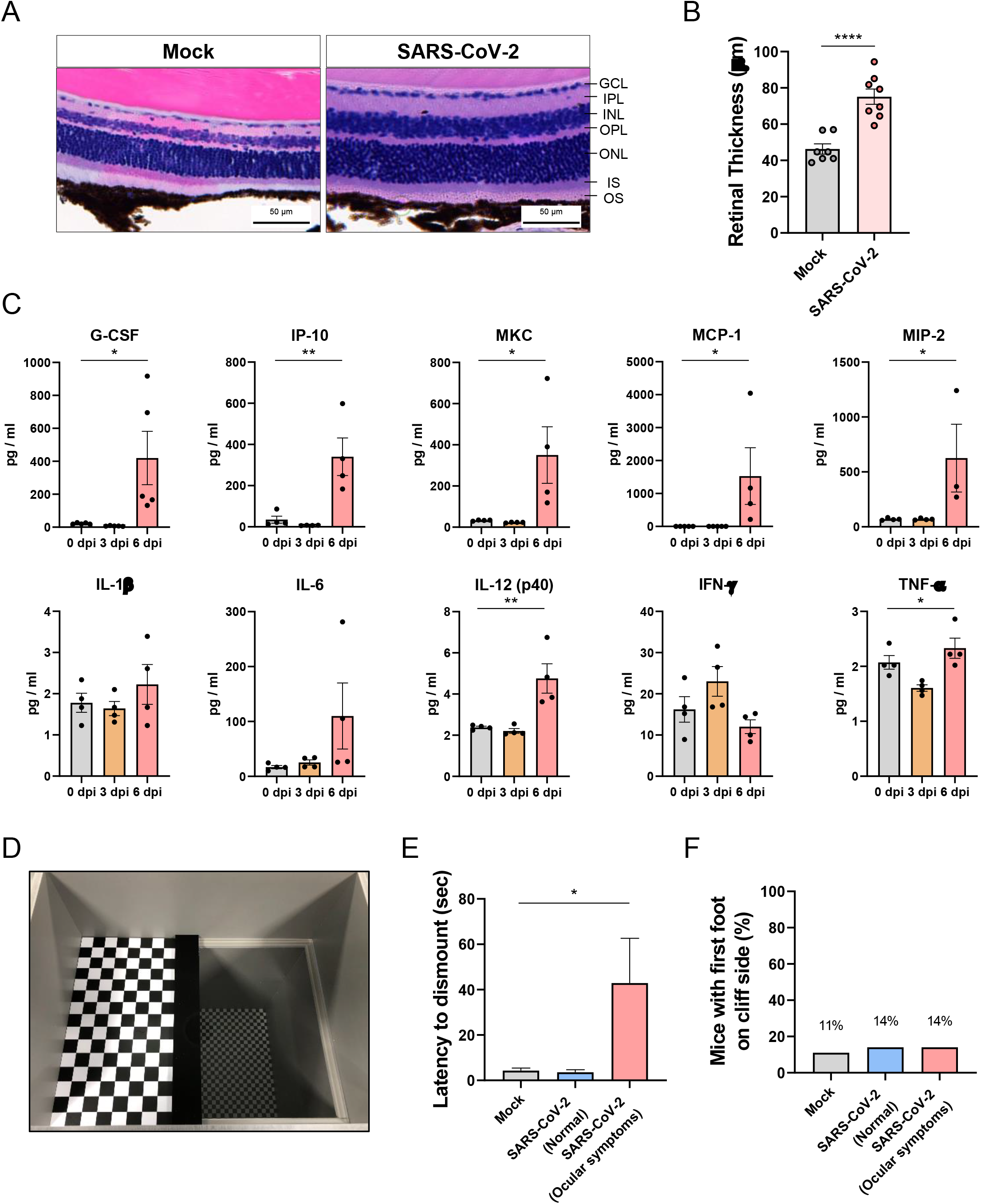
Histopathological and multiplex cytokine analyses of the eyes of SARS-Cov-2-infected mice. H&E staining of the eye sections from K18-hACE2 mice six days post mock infection or SARS-CoV-2 infection. **(A)** Representative histological images (*n* = 4 per group) show changes in the retinal thickness. Scale bar = 50 μm. **(B)** The retinal thickness was measured and shown in a bar graph (Mock, Gray; SARS-CoV-2, Red). **(C)** The chemokine and cytokine levels of eyes were measured by multiplex immuno-analysis (*n* = 4 per indicated dpi; 0 dpi, Gray; 3 dpi, Yellow; 6 dpi, Red). G-CSF, Granulocyte–macrophage colony-stimulating factor; IP-10, C-X-C motif chemokine 10 (CXCL10); MKC, mouse keratinocyte-derived chemokine; MCP-1, Monocyte Chemoattractant Protein-1 (CCL2); MIP-2, Macrophage-inflammatory protein 2 (CXCL2); **(D)** The photo of the visual cliff apparatus used in this study. **(E-F)** Time for latency to dismount platform (E) and the percent of mice that took the first footstep over the cliff side (F) were measured (*n* ≥ 7 per groups). Symbols represent means ± SEM. Statistically significant differences between the groups were determined by Student’s t-test (B) or one-way ANOVA (C, E). *\*P* < 0.05; \*\**P* < 0.01; \*\*\*\**P* < 0.0001. SARS-CoV-2: severe acute respiratory syndrome coronavirus 2; GCL: ganglion cell layer; IPL: inner plexiform layer; INL: inner nuclear layer; OPL: outer plexiform layer; ONL: outer nuclear layer; IS: inner segments; OS: outer segments; dpi: days post-infection

Next, we evaluated the functional consequences of the retinal inflammation on visual behavior. The innate aversion to depth has been exploited in the visual cliff test designed to study depth perception in mice using a visual cliff apparatus shown in Fig. 2D (Fox, 1965). The test order was randomized and each mouse was tested only once in a life time to avoid a memory effect. SARS-CoV-2-infected mice were divided into two groups on 5 dpi according to the presence or absence of ocular symptoms. On the same day, the mice of mock and SARS-CoV-2-infected groups dismounted the platform within 4.28 and 3.56 s (mean), respectively (Fig. 2E). SARS-CoV-2-infected mice with ocular symptoms presented a prolonged latency for the dismounting (42.92 sec). However, there was no difference in the number of mice with first foot on the cliff side between mock and SARS-CoV-2-infected groups (Fig. 2F), indicating that the retinal inflammation induced by viral infection did not exacerbate to retinal degeneration or visual loss.

### Ocular tropism of SARS-CoV-2 through neuronal invasion in mice

Similar to other neurotropic viruses such as ZIKV and West Nile virus, SARS-CoV and SARS-CoV-2 have been reported to cause symptoms of neurological disorders in patients (Ramani et al., 2021; Zhou et al., 2020b). Recently, evidence of rapid SARS-CoV-2 spread to the olfactory bulb via olfactory nerves was found in patients with COVID-19 (de Melo et al., 2021; Meinhardt et al., 2021). Since both the olfactory nerve and TN provide anatomical connections between the brain and nasal passages (Lochhead and Davis, 2019), we hypothesize that IN-administered SARS-CoV-2 spreads to the brain and eyes via the TN and ON (Fig. 3A). To validate this hypothesis, we infected mice with SARS-CoV-2 using diverse injection routes (Fig. 3B; IT: intratracheal, IC: intracerebral, ED: eye-drop, IV: intravenal). Approximately 10^4^ PFU was injected into the mice. We then assessed viral loads in the lungs, brain, eyes, TN, and ON on 3 and 6 dpi. An overall weight loss in mice injected via the IN, IT, and IC routes was observed. Mortality was observed only in mice injected via the IC route on 2 dpi (Supplementary Fig. 3). This result is consistent with that of a previous study that demonstrated that brain infection of SARS-CoV leads to death in K18-hACE2 mice (Netland et al., 2008). The IN injection resulted in ocular tropism with increased viral RNA levels in the eyes between 3 and 6 dpi (Fig. 3C). In general, the viral RNA titers detected in the TN and ON were almost equal to those in the brain and eyes, indicating viral transmission to the eyes and brain via the neuronal invasion of these nerves. Yet, since the nasolacrimal duct provides an anatomical connection between the ocular surface and respiratory tract (Willcox et al., 2020), the viral spread to the eyes could have unintentionally occurred via the nasolacrimal duct, and not via the neurons. To rule out this, we infected via the IT route. In a similar manner to IN infection, increased viral RNA titers were also detected in the eyes, brain, TN, and ON on 6 dpi, indicating viral transmission from the lung to the eyes and brain through TN and ON. To confirm the infection route between the eyes and brain through TN and ON, we directly administered the virus into the brain through an IC injection. The robust viral RNA copies were found in the eyes, brain, TN, and ON, but were relatively lower than those in the lung. IC infection evidently showed that the virus in the brain can migrate to eyes through the TN and ON. In addition, as both IN and IC infections showed the highest viral titer in the eyes on 6 dpi, tearing and eye discharge were found only in mice infected via the IN and IC routes. These results revealed that viral transmission occurs between the brain and eyes through the TN and ON, with a network reaching the lung.

**Figure 3.**
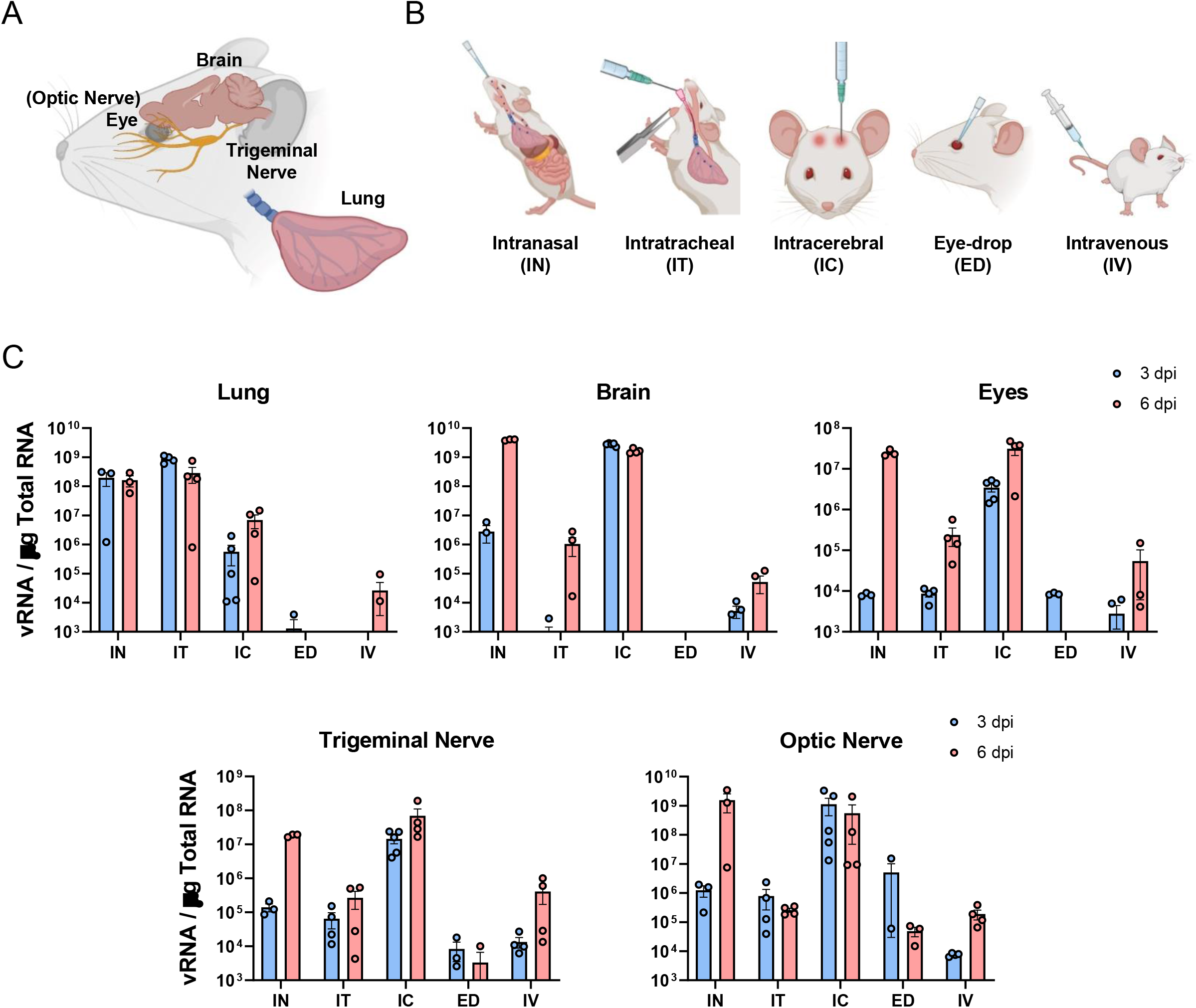
Virus titers of SARS-CoV-2-infected mice via diverse injection routes. **(A)** An anatomical graphic for the visual depiction of murine brain, eye (optic nerve), trigeminal nerve, and lung. **(B)** Illustrations for diverse injection routes: Intranasal (IN), Intratracheal (IT), Intracerebral (IC), Eye-drop (ED), and Intravenous (IV) routes. K18-ACE2 mice were inoculated with approximately 10^4^ PFU SARS-CoV-2 via five different injection routes (n = 8 per injection route; n = 4 for 3 dpi and 6 dpi, respectively). **(C)** Viral RNA levels in the lungs, brain, eyes, trigeminal nerve, and optic nerve were analyzed on 3 and 6 dpi by RT-qPCR (3 dpi, Blue; 6 dpi, Red). Viral RNA copies were cut-off by the limit of detection (10^3^ copies/μg). Symbols represent means ± SEM. SARS-CoV-2: severe acute respiratory syndrome coronavirus 2; PFU: plaque-forming unit; dpi: days post-infection

The ocular surface is considered as an additional mucosal surface exposed to virus-containing aerosols. To investigate whether the virus can enter the eyes and migrate to the respiratory tract, we measured the viral load following the ED infection (approximately 10^4^ PFU). The viral RNA load was detected in the eyes on 3 dpi but disappeared on 6 dpi, and low viral load was detected only in the TN and ON. This result demonstrated the faint possibility of ocular susceptibility to SARS-CoV-2. A previous study suggested that SARS-CoV-2 infection by the IV route was not required for the lung infection (Hassan et al., 2020). Indeed, the IV infection resulted in relatively low viral copies in every tissue studied; the lungs, brain, eyes, TN, and ON. Overall, SARS-CoV-2 can transmit from the respiratory tract to eyes through the brain tracking of TN and ON, which cannot be reversed.

### Ocular- and neuro-tropism of infectious SARS-CoV-2-mCherry in mice

The incorporation of a fluorescent reporter gene into a replication-competent virus has advanced our ability to trace viral infection and tropism in vivo. To confirm the viral spread from the respiratory tract to eyes through the brain, we used an infectious SARS-CoV-2-mCherry clone generated by the simple manipulation of the reverse genetics system (Rihn et al., 2021). A sequence encoding the mCherry fluorescent marker was inserted in-frame after the C-terminus of the ORF7a protein to avoid deletion of viral sequences. This virus can replicate and recapitulate features of severe COVID-19 infections associated with the mice model (Liu et al., 2021b). In mice infected via the IN route with approximately 10^4^ PFU of SARS-CoV-2-mCherry clone, significant morbidity (Fig. 4A) and mortality (Fig. 4B) were observed from 7 dpi. The viral distribution in tissues including the lungs, brain, eyes, spleen, TN, and ON was analyzed on 6 dpi by an *in vivo* imaging system (IVIS) to detect the fluorescence of SARS-CoV-2-mCherry. For the fluorescence detection of the infection in the TN and ON of mice, we cut the exposed skull along the left and right sides, to between the eyes, and then pulled the brain out, exposing the base of the cranial cavity (Fig. 4C). Robust fluorescence signals were detected in every tissue studied (ranged from 10^6^ to 10^9^ radiant efficiency [p/s/cm^2^/sr]/[μW/cm^2^]) and were clearly distinguishable from that of mock group except for the spleen that served as a negative control (Fig. 4D). The viral distribution to the brain and eyes validated the neuronal invasion of the virus to the TN and ON that can be used for SARS-CoV-2 transmission.

**Figure 4.**
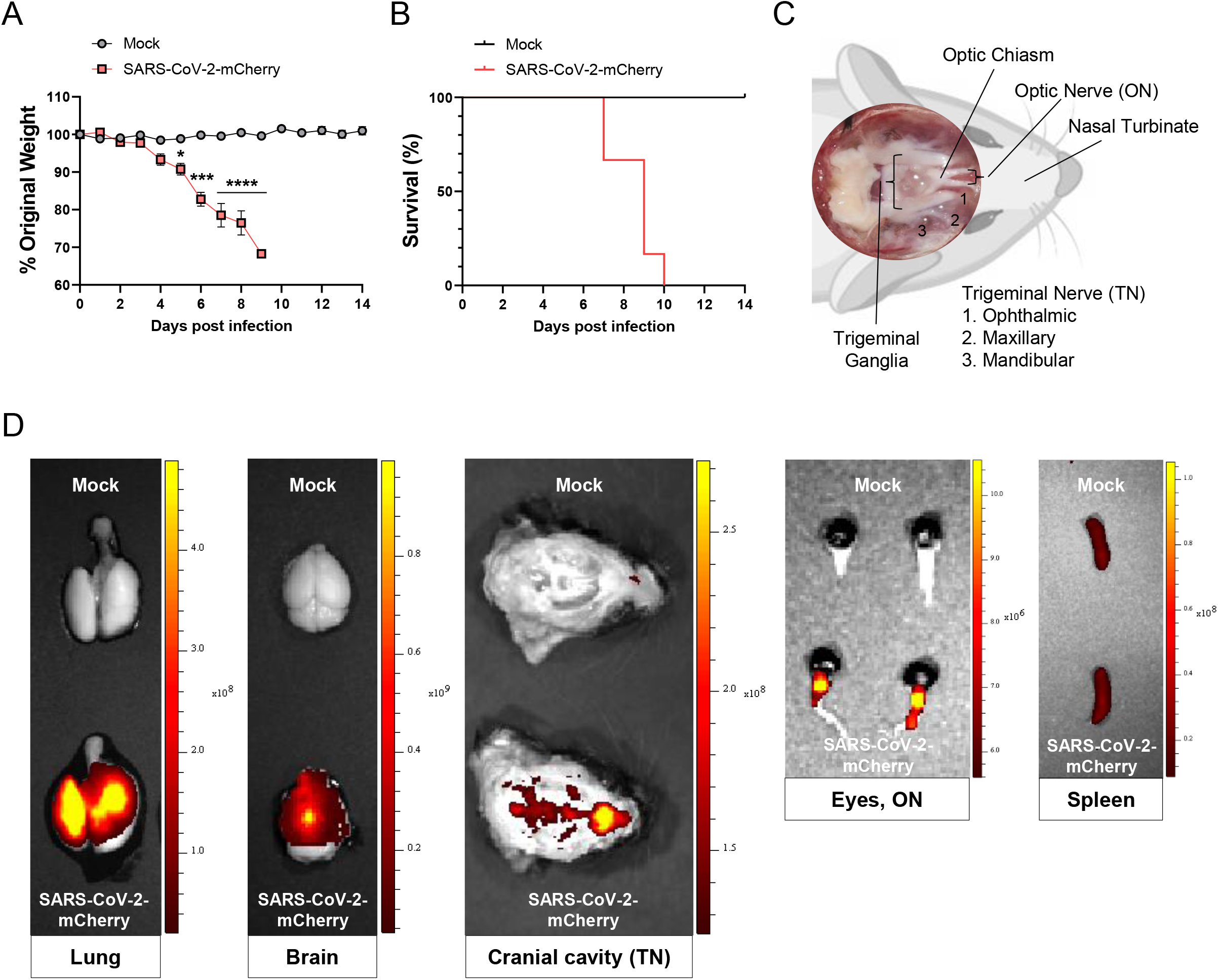
*In vivo* viral distributions of infectious SARS-CoV-2-mCherry in IN-infected mice. K18-hACE2 mice were IN-inoculated with approximately 10^4^ PFU of SARS-CoV-2-mCherry (*n* = 6 for the mock-infected and infected mice, respectively). **(A)** Body weight and **(B)** survival were monitored on the indicated dpi. Symbols represent means ± SEM. Statistically significant differences between the groups were determined by multiple t-test. \**P* < 0.05; \*\*\**P* < 0.001; \*\*\*\**P* < 0.0001. **(C)** A brain-dissected dorsal view of the murine cranial cavity showing optic nerves, optic chiasm, trigeminal ganglia, and trigeminal branches (1, Ophthalmic; 2, Maxillary; 3, Mandibular). **(D)** Representative *in vivo* fluorescence images of organs, including the lungs, brain, eyes, and spleen in mock- or SARS-CoV-2-mCherry-infected mice (*n* = 5 for the mock-infected and infected mice, respectively; Upper, Mock; Lower, SARS-CoV-2-mCherry) were acquired by sequential imaging. Color bars indicate radiant efficiency [p/s/cm^2^/sr]/[μW/cm^2^]. SARS-CoV-2: severe acute respiratory syndrome coronavirus 2; PFU: plaque-forming unit; dpi: days post-infection

### Ocular tropism of SARS-CoV-2 through the brain via the TN and ON in wild-type Syrian hamsters

In K18-hACE2 mice, since the expression and distribution of hACE2 are under the control of the cytokeratin 18 promoter (McCray Jr et al., 2007), they are not naturally sensitive to SARS-CoV-2 infection. Investigation of the ocular tropism in wild-type Syrian hamsters, of which an endogenous ACE2 protein can interact with SARS-CoV-2 to make it naturally permissive to viral infection (Chan et al., 2020; Imai et al., 2020; Luan et al., 2020; Sia et al., 2020), needs to be extrapolated to humans. SARS-CoV-2 IN-inoculation (approximately 10^4^ PFU) of Syrian hamsters resulted in weight loss (~ 10%) on 6 dpi, but animals almost returned to their original weight by 14 dpi (Fig. 5A). During the course of the experiments, no mortality was observed in both mock and infection groups. To determine the ocular tropism of SARS-CoV-2 in a hamster model, we infected the virus via IN and ED routes. Compared to that with IN infection, there was no significant weight loss for ED-infected (approximately 10^4^ PFU) hamsters (Fig. 5B). At necropsy, gross pathology revealed pathological lesions and lung congestion only in IN-infected hamsters on 6 dpi (Fig. 5C). The quantity of infectious SARS-CoV-2 in the lungs along with the eyes at 6 days after IN or ED infection was assessed. The presence of infectious virus in eyes was found in 60% of IN-infected hamsters, despite lower viral titers (mean, 2.5 × 10^1^ PFU/ml) than that in the lungs (mean, 1.6 × 10^3^ PFU/ml), confirming the ocular tropism of SARS-CoV-2 in wild-type animals (Fig. 5D, 5E). Further virus titer analysis by RT-qPCR showed the presence of virus in other tissues of IN-infected hamsters, including the brain, ON, and TN at higher titers than that in spleen, which is barely susceptible to SARS-CoV-2 infection (Fig. 5F). However, marginally higher viral RNA levels were detected only in the eyes and ON of ED-infected hamsters compared to those in the spleen (Fig. 5G). These data are consistent with the aforementioned findings in K18-hACE2 mice suggesting that SARS-CoV-2 can spread from the respiratory tract to the brain and eyes via the TN and ON, which cannot be reversed.

**Figure 5.**
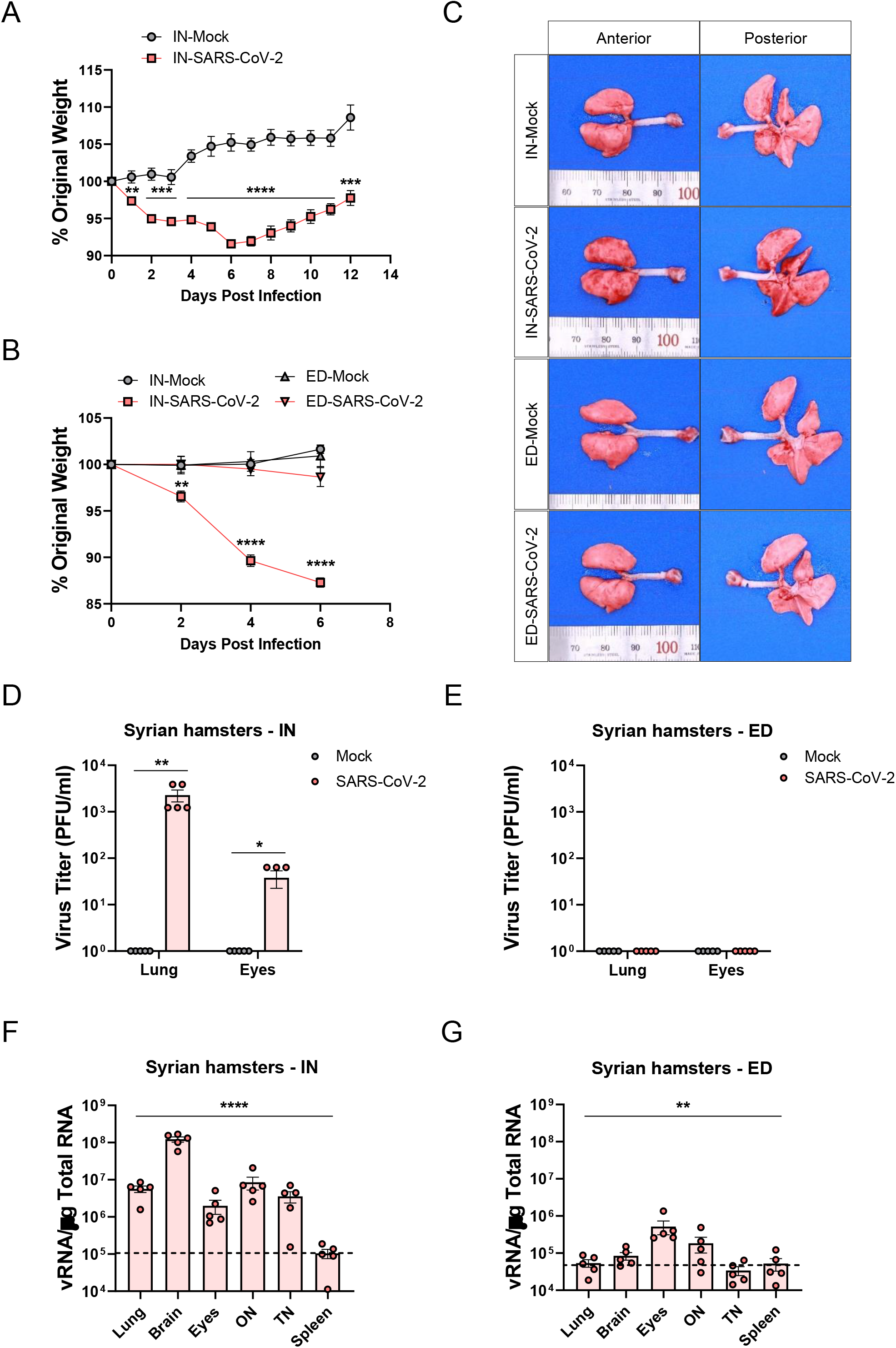
Clinical features, gross pathology, and viral titers of SARS-CoV-2 in intranasally (IN) or eye-drop (ED)-infected wild-type Syrian hamsters. Eleven-week-old female Golden Syrian hamsters were IN-mock-infected or infected with approximately 10^4^ PFU of SARS-CoV-2 (*n* = 6 for mock-infected and *n* = 5 for infected mice; mock, gray; SARS-CoV-2, red). **(A)** Body weight changes are shown as a percentage of the starting weight at the indicated dpi after IN infection. **(B)** Syrian hamsters were IN- or ED-infected with approximately 10^4^ PFU of SARS-CoV-2 (each *n* = 5 for IN-mock-infected, IN-infected, ED-mock-infected, and ED-infected). A graph showing the percent body weight change is shown. **(C)** Representative digital images of lungs harvested from mock-infected or infected hamsters on 6 dpi. **(D-E)** Viral loads in the lungs and eyes of IN-infected (D) and ED-infected (E) hamsters via the median tissue culture infectious dose (TCID_50_) assay. **(F-G)** Viral RNA levels in the lungs, brain, eyes, optic nerves (ONs), trigeminal nerves (TNs), and spleens of IN-infected (F) and ED-infected (G) animals were assessed by RT-qPCR. Viral RNA copies were subjected to a cut-off based on the limit of detection (10^4^ copies/μg). A dashed line indicates the viral RNA levels in the spleen, which is barely susceptible to SARS-CoV-2 infection. Symbols represent means ± SEMs. Statistically significant differences between the groups were determined by a multiple t-test (A, B), Student’s t-test (D, E), or one-way ANOVA (F, G). \**P* < 0.05; \*\**P* < 0.01; \*\*\**P* < 0.001; \*\*\*\**P* < 0.0001. SARS-CoV-2: severe acute respiratory syndrome coronavirus 2; PFU: plaque-forming unit; dpi: days post-infection

## Discussion

A previous study highlighted SARS-CoV-2 neuro-tropism and suggested the need to identify the route of viral invasion into the brain (Song et al., 2021). Olfactory nerves form bundles that provide an anatomical connection between the brain and nasal passage through foramina in the cribriform plate and synapse on glomeruli in the olfactory bulb (Norwood et al., 2019). In addition, branches V_1_ (ophthalmic) and V_2_ (maxillary) of TN (Fig. 4C) innervate both the respiratory and olfactory regions of the nasal passage, connecting the brain (Bojsen□Møller, 1975; Schaefer et al., 2002). Although SARS-CoV-2 infection and transmission of olfactory nerves have been demonstrated by several groups (de Melo et al., 2021; Meinhardt et al., 2021; Zhang et al., 2021), those of TNs remain to be elucidated. Neuronal invasion via TNs has been suggested in humans, based on a patient with COVID-19 who suffered from trigeminal neuralgia (Molina-Gil et al., 2021). In this study, we demonstrated that TNs can be infected by SARS-CoV-2 and are used for the viral transmission to the brain and eyes, along with the ONs.

Subtype H7 influenza viruses have the ocular tropism and use the eyes as an entry route, which was confirmed by dropping the virus onto the corneal surface (Belser et al., 2013). Similarly, through eye-dropping of SARS-CoV-2, we certainly excluded the possibility of ocular surface as a portal of entry for the virus (Fig. 3B, 5E, 5F). The viral burdens of the eyes were comparatively low and disappeared as time passed (Fig. 3B). This may be due to the tear film and blinking. There are anti-viral factors in the tear film, such as lysozyme, lipocalin, and lactoferrin that provide an immunological and protective environment against viral infections. The act of blinking provides not only physical protection from outside contaminants but also helps in the drainage of the tear film from the tear punctum (de Freitas Santoro et al., 2021).

Clinical studies have reported that the time required for ocular manifestation varied from 15 days to two months after the infection or symptom onset (Gasparini et al., 2021; Yan et al., 2020). Moreover, Colavita et al detected viral RNA in ocular swabs with lower Ct values than those in nasal swabs between 21 and 27 days from the symptom onset of a patient with COVID-19 who suffered from ocular manifestations (Colavita et al., 2020). Interestingly, they also isolated live replication-competent viruses directly from the ocular fluid collected from the patient. Consistent with these clinical studies, IN- and IT-administered viral copies in eyes increased in a time-dependent manner (Fig. 3C).

In summary, ocular manifestation and retinal inflammation were promoted by SARS-CoV-2 infection in the mouse model, inducing the increased cytokine production. The virus spreads from the lungs to the brain and eyes through the network consisting of TN and ON. This ocular tropism was also observed in wild-type Syrian hamsters. It would be interesting and important to further investigate how the viral infection of eyes elicits ocular inflammation and manifestations and their clinical relevance. Along with the respiratory system, eyes and TNs should be considered SARS-CoV-2-susceptible organ systems. Our data raise awareness of ocular and neuronal infection-mediated disorders beyond respiratory diseases to set up treatment strategies for patients with COVID-19.

## Materials and methods

All procedures were performed in a biosafety level 3 (BSL3) or animal BSL3 facility for SARS-CoV-2-related experiments, with approval from the Korea Research Institute of Chemical Technology (KRICT) and by personnel equipped with powered air-purifying respirators.

### Animals

#### Mice

Eight-week-old male B6.Cg-Tg(K18-hACE2)2Prlmn/J mice were purchased from the Jackson Laboratory and maintained in a biosafety level 2 (BSL-2) animal facility in the Korea Research Institute of Chemical Technology (KRICT). All protocols were approved by the Institutional Animal Care and Use Committee (Protocol ID 8A-M6, IACUC ID 2021-8A-02-01 & 2021-8A-03-03).

#### Syrian hamsters

Eleven-week-old female Golden Syrian hamsters were purchased from Janvier Labs (Saint-Berthevin, France). The hamsters were kept in standard cages and exposed to a 12:12 hour light/dark cycle at 22–24°C, with 40–55% humidity and food and water supplied *ad libitum*, in an animal biosafety level 3 (ABL-3) facility in the Korea Research Institute of Bioscience & Biotechnology (KRIBB). All protocols were approved by the Institutional Animal Care and Use Committee (KRIBB-ACE-21329, KRIBB-IBC-20220201).

### Cells and Viruses

The SARS-CoV-2 Korean strain (GISAID Accession ID:EPI_ISL_407193), isolated from a Korean patient, was obtained from the Korea Disease Control and Prevention Agency (KCDC) and propagated in Vero cells (CCL-81, American Type Culture Collection (ATCC), Manassas, VA, USA). pCC1-4K-SARS-CoV-2-mCherry clone (GenBank Accession No. MT926411) was used for the rescue of infectious virus following the protocol developed by Rihn et al (Rihn et al., 2021). Briefly, 3 μg of plasmid DNA was transfected into BHK-21 cells (CCL-10, ATCC) in a six-well plate using Lipofectamine LTX with Plus reagent (15338100, Invitrogen, Waltham, MA, USA) as per the manufacturer’s instructions. Three days post transfection, the supernatant was transferred to Vero cells in a T25 flask. After further incubation for four days, the infectious virus was titrated by a plaque assay.

### Virus titer

#### Plaque assay

The virus was serially diluted in Eagle’s minimum essential medium supplemented with 2% fetal bovine serum for the plaque assay. Culture medium was removed from 24-well plated Vero E6 cells (approximately 1 × 10^5^ per/well) a day before the assay, and the inoculum was transferred onto triplicate cell monolayers. After incubation at 37 °C for 1 h, the inoculum was removed, and infected cells were overlaid with 1.8% carboxymethyl cellulose in MEM. Samples were incubated for four days, followed by fixation and staining with 0.05% crystal violet containing 1% formaldehyde. The counts of plaques were measured by an ImmunoSpot analyser (Cellular Technology Ltd, Shaker Heights, OH, USA).

#### Median tissue culture infectious dose (TCID_50_) assay

To determine the titer of SARS-CoV-2, TCID_50_ assays were performed with Vero cells, as described elsewhere (Sia et al., 2020). Ten-fold serial dilutions of stock viruses were prepared in DMEM. From these dilutions, 100 μl each was transferred to monolayers of Vero cells grown in 96-well plates in DMEM supplemented with 1 μg/ml TPCK-treated trypsin and incubated at 37°C in a 5% CO_2_ incubator. Virus titers were calculated at 4 days post-infection and expressed as TCID50/ml values based on the method of Reed and Muench (Reed & Muench, 1938). The virus dilutions were made based on a prediction of the number of plaque forming units (PFU) using a conversion formula as follows: PFU (ml) / TCID50 (ml) = 0.7 (Davis et al., 1972).

### Viral inoculations

All viral inoculations (SARS-CoV-2 diluted in PBS, approximately 10^4^ PFU) were administered by IN, IT, IC, ED, and IV routes under anesthesia using isoflurane in a BSL-3 animal facility, and all efforts were made to minimize animal suffering. IN, IC, and IV injections were performed as per the protocol of a previous study (Shimizu, 2004). Mock group was injected with the same volume of PBS in all the experiments. Body weights were measured everyday post-infection.

#### IN injection

Mice were infected with a volume of 20 μl per mouse. The mouse was anesthetized and the inoculum was administered dropwise into one nostril.

#### IT injection

For the IT injection, the protocol of a previous study was followed (DuPage et al., 2009). In brief, mice were injected with a volume of 20 μl per mouse. The anesthetized mice were placed on the string by their front teeth, with vertical hanging of their chest on the platform. A high intensity of light illuminated the upper chest. The mouth was opened, and the tongue was pulled out using the flat forceps to see the white light from the trachea. The catheter was inserted into the trachea, and then, the needle was removed. The inoculum (20 μl) was directly injected into the opening of the catheter. The IT injection was performed with 2% solution of Evans Blue in normal saline to rehearse this injection (E2129, Sigma-Aldrich, St. Louis, MO, USA; Supplementary Fig. 4)

#### IC injection

The anesthetized mice were injected with a volume of 10 μl per mouse. Before the injection, the inoculum was loaded into the glass microliter syringe (80401, Hamilton, Reno, NV, USA) with a 30 G × 4 mm ultra-fine disposable needle (0J293, Jeongrim medical, Chungcheongbuk-do, Republic of Korea). The injection site was half way between the eyes and ears and just off the midline. The needle was used to directly and slowly penetrate the cranium. The injection was performed very slowly and was followed by a very slow removal to prevent efflux. The IC injection was performed with 2% solution of Evans Blue in normal saline to rehearse this injection (Supplementary Fig. 4).

#### ED

The anesthetized mice were injected with a volume of 4 μl per eye. The inoculum was administered dropwise onto the corneal surface, followed by massaging the eyelids.

#### IV injection

Mice were infected with a volume of 50 μl per mouse. They were placed in the tail access rodent restrainers. The inoculum was slowly injected into the tail vein.

### Quantitative RT-PCR

The total RNA of tissues was extracted using the Maxwell RSC simply RNA tissue kit (AS1340, Promega, Madison, WI, USA) following the manufacturer’s protocol. Quantitative RT-PCR (QuantStudio 3, Applied Biosystems, Foster City, CA, USA) was performed using one-step Prime script III RT-qPCR mix (RR600A, Takara, Kyoto, Japan). The viral RNA of nucleocapsid protein was detected by a 2019-nCoV-N1 probe (10006770, Integrated DNA Technologies, Coralville, IA, USA).

### Multiplex analysis

The eyes of SARS-CoV-2-infected mice were dissected on 0, 3, and 6 dpi, and then homogenized in bead tubes (a-psbt, GeneReach Biotechnology, Taichung, Taiwan). Aliquots of them were analyzed with the MILLIPLEX human cytokine/chemokine magnetic bead panel (HCYTOMAG-60K, Merck Millipore, Burlington, MA, USA) using the Luminex 200 multiplexing instruments (40-012, Merck Millipore) to assess the cytokine/chemokine expression.

### H&E staining

The eyes were collected and fixed in Davidson’s fixative (BBC Biochemical, Mount Vernon, WA, USA) overnight. The fixed eyes were processed routinely and embedded in paraffin wax (ASP300S, Leica, Wetzlar, Germany). After embedding, the eyes were cut into 4 μm sections and stained with H&E (BBC Biochemical) using an autostainer (ST5010, Leica). Images were obtained using an Olympus BX51 microscope (Olympus, Tokyo, Japan). The retinal thickness was measured using Nuance 3.02 software (PerkinElmer, Waltham, MA, USA).

### Visual cliff test

Visual cliff test was carried out based on the protocol developed by Tzameret et al (Tzameret et al., 2015) with some modifications. The mice were tested in an open-topped acrylic glass box (a dimension of 40 × 30 × 50 cm). The light source in the experimental room was dimmed (about 20 lx). The barriers were opaque to prevent reflection. A paper with a large black/white checkered pattern (2 × 2 cm square) was placed under the half of plate (‘bench side’) while the bottom of the other half (‘cliff side’) was covered by a small checkered pattern sheet (1 × 1 cm square) to emphasize the cliff drop-off. The mice were placed on the black central platform (0.4 × 30 × 0.1 cm) between the bench side and the cliff side, and their activity was recorded for 2 min to measure the latency to dismount and the direction of the first foot on the bench or cliff sides.

### *In vivo* fluorescence spectrum unmixing

SARS-CoV-2-mCherry-infected mice were sacrificed on 6 dpi, and perfused with 4% paraformaldehyde in PBS. The lungs, brain, and other organs were immediately collected to detect fluorescent signals. Using IVIS Lumina S5 system (PerkinElmer, Wrentham, MA, USA), the tissues were sequentially imaged with a fixed emission filter at 520 nm to determine the optimal excitation in the range from 420 nm to 480 nm (1 sec, F-stop = 1, medium binning). Images were analyzed using Living Image 4.7 software (PerkinElmer, Wrentham, MA, USA). The epi-fluorescence was expressed in units of radiant efficiency [p/s/cm^2^/sr]/[μW/cm^2^] after subtracting the background signal.

### Statistical analysis

All experiments were performed at least three times. All data were analyzed using the GraphPad Prism 8.0 software (GraphPad Software, San Diego, CA, USA). Data are shown as mean with the SEM. For experiments with only two groups, Student’s t-test (two-tailed) was used. For groups with three and more groups, one-way analysis of variance (ANOVA) was used. *P* < 0.05 was considered statistically significant. Specific analysis methods are described in the figure legends. Statistical significance is indicated as follows: * *P* < 0.05; ** *P* < 0.01; *** *P* < 0.001, **** *P* < 0.0001.

## Online supplemental material

Fig. S1 shows the up-regulated cytokines in the brains of SARS-CoV-2-infected mice using the multiplex analysis. Fig. S2 shows the results of IT and IC infections with Evans blue. Fig S3 indicates the body weight loss and mortality by diversely administered SARS-CoV-2 infections in the mouse model.

## Abbreviations

COVID-19: coronavirus disease 2019
SARS-CoV-2: severe acute respiratory syndrome coronavirus 2
PFU: plaque-forming unit
IN: Intranasal
IT: Intratracheal
IC: Intracerebral
ED: Eye-drop
IV: Intravenous
TN: trigeminal nerves
ON: optic nerve

## Author contributions

Conceptualization: G.U.J., and Y.-C.K.; Methodology: G.U.J., H.-J.K. H.W.M., and Y.-C.K.; Investigation: G.U.J., H.-J.K., H.W.M., G.Y.Y., H.-J.S., J.S.C., I.-C.L., and D.-G.A.; Resources: H.-J.K., S.-J.K. S.M., and Y.-C.K.; Data curation and analysis: G.U.J. and Y.-C.K.; Writing–original draft: G.U.J., and H-J.K.; Writing–review and editing: G.U.J., S.-J.K., K.-D.K., S.M. and Y.-C.K; Visualization: G.U.J. and Y.-C.K.; Supervision: Y.-C.K.; Funding acquisition: H.-J.K. and Y.-C.K.

## Acknowledgements

Graphic abstract and other incorporating images in figures were created using BioRender.

This work was supported by grants from the National Research Council of Science & Technology (NST) funded by the Korean government (MSIP; CRC-16-01-KRICT to Y.-C.K.), (MSIT; CRC21021 to H.-J.K.) and the National Research Foundation of Korea (NRF) funded by the Ministry of Education, Science, and Technology (MSIT) of the Korean government (2020R1C1C1003379 to Y.-C.K.).

The authors declare no competing financial interests.

## Figure legends

**Figure S1.**
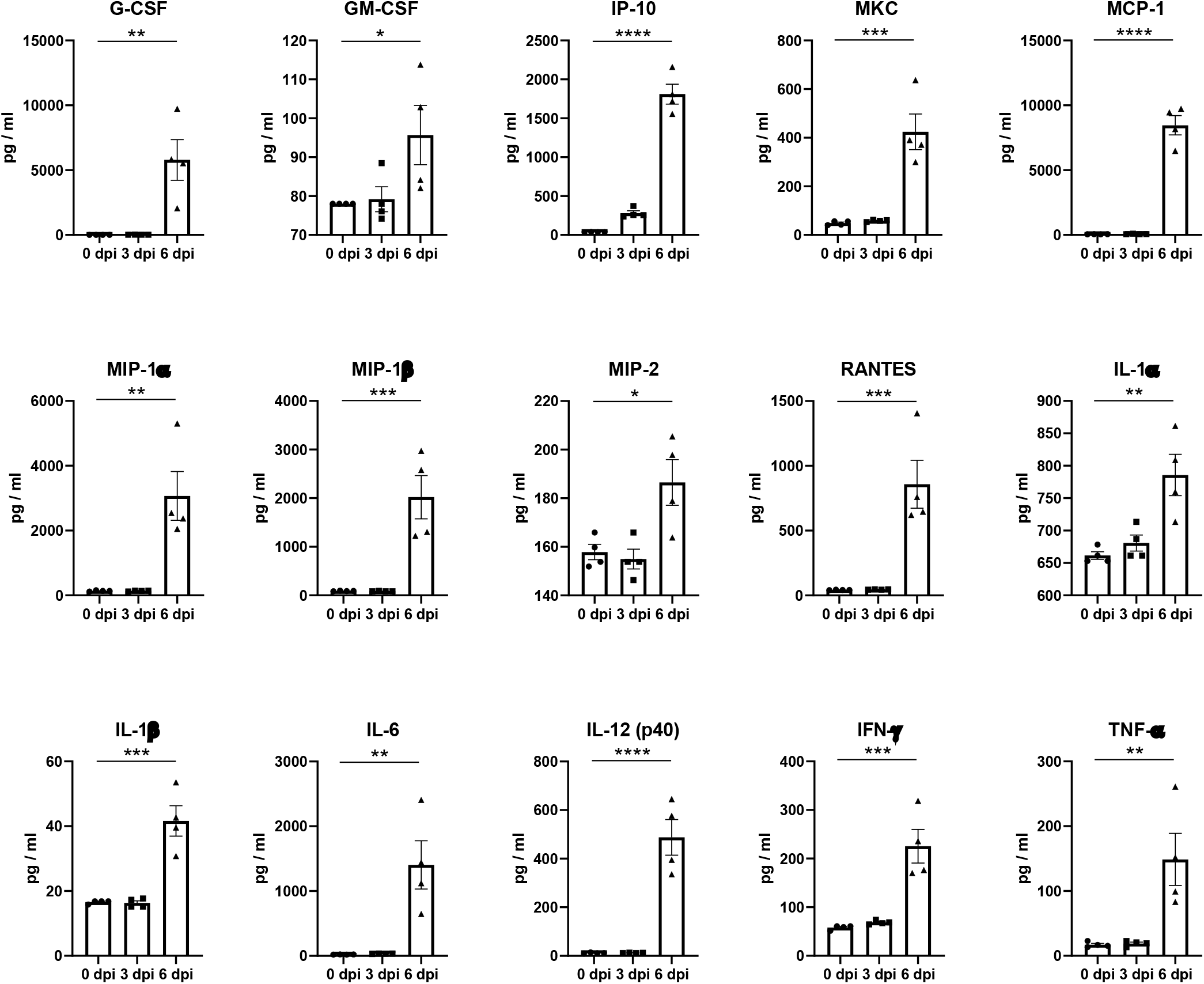
Multiplex cytokine analysis of the brain tissues of SARS-CoV-2-infected mice. The chemokine and cytokine levels of the brain were measured by multiplex immuno-analysis (*n* = 4 per indicated dpi). G-CSF, Granulocyte–macrophage colony-stimulating factor; IP-10, C-X-C motif chemokine 10 (CXCL10); MKC, mouse keratinocyte-derived chemokine; MCP-1, Monocyte Chemoattractant Protein-1 (CCL2); MIP, Macrophage-inflammatory protein; RANTES, Regulated upon Activation, Normal T Cell Expressed and Presumably Secreted (CCL5); Symbols represent means ± SEM. Statistically significant differences between the groups were determined by one-way ANOVA. \**P* < 0.05; \*\**P* < 0.01; \*\*\**P* < 0.001; \*\*\*\**P* < 0.0001. SARS-CoV-2: severe acute respiratory syndrome coronavirus 2; dpi: days post-infection

**Figure S2.**
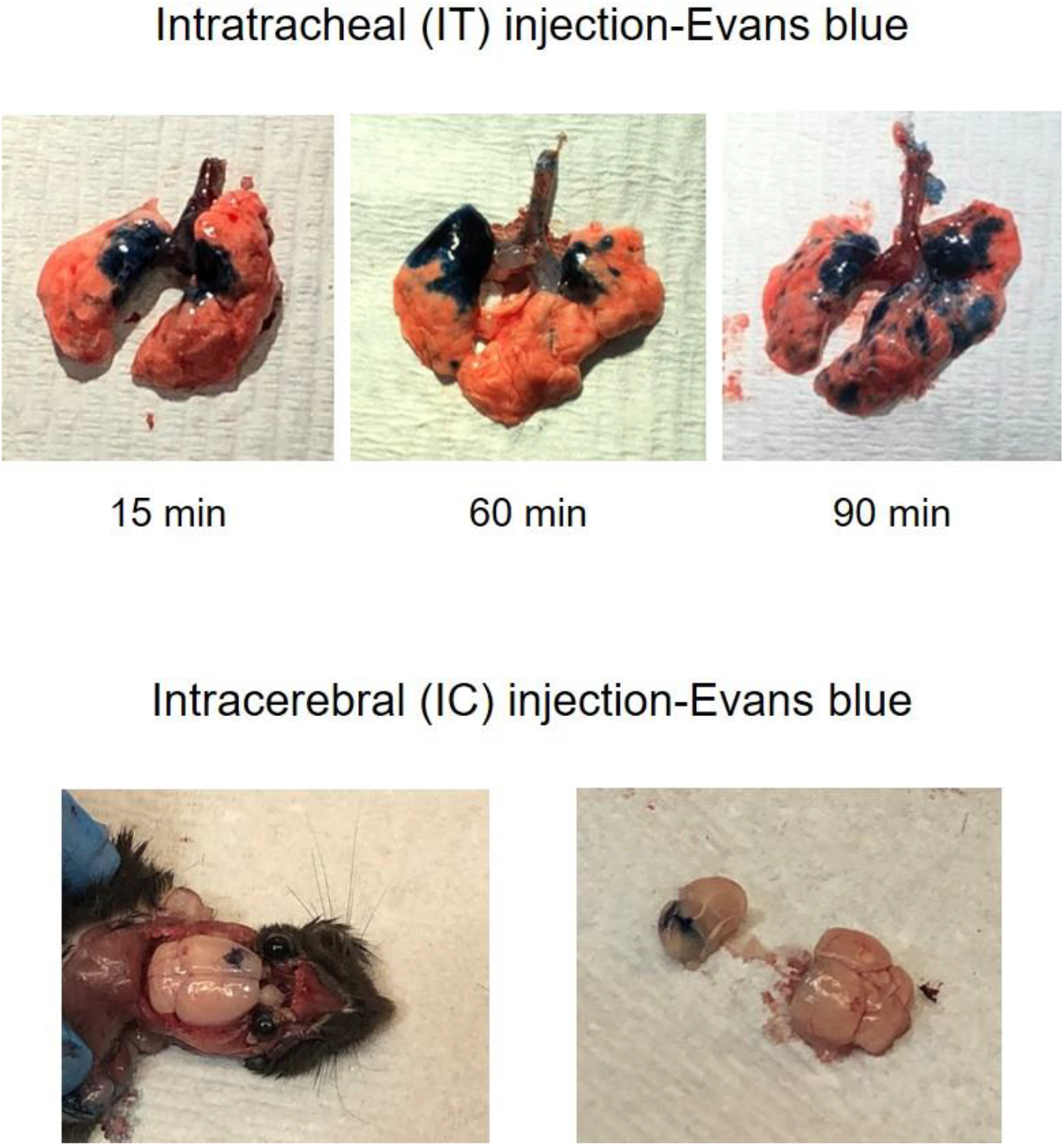
Intratracheal and intracerebral injection of Evans blue. Images of the lung (Upper) at indicated time and brain (Lower) following intratracheal and intracerebral injections with 2% solution of Evans Blue in normal saline.

**Figure S3.**
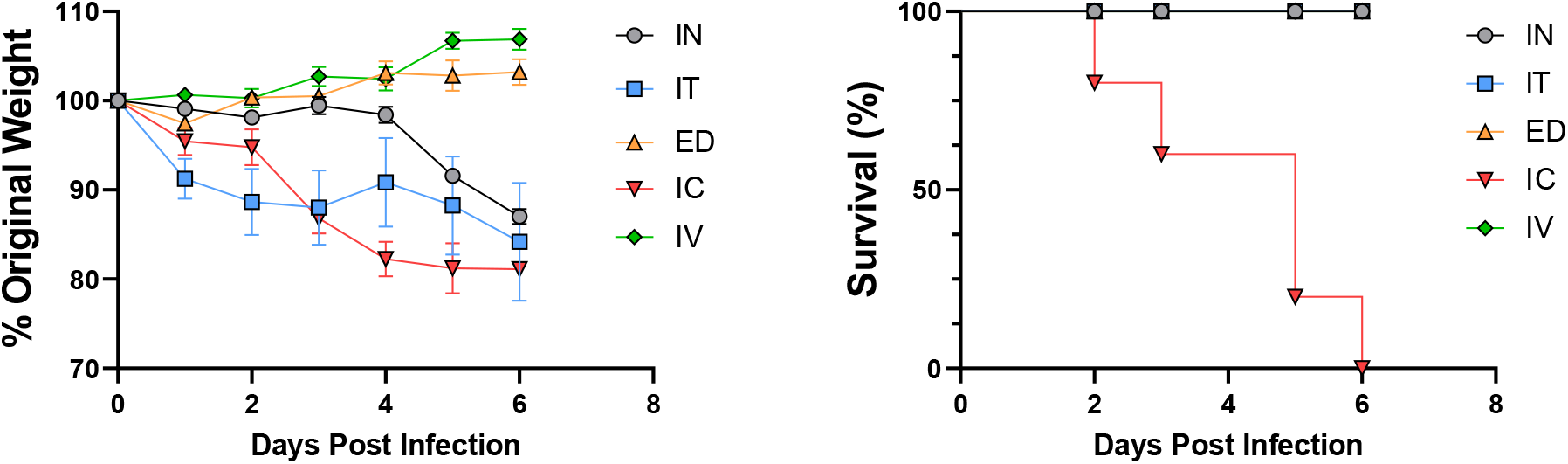
Body weight and survival rate following SARS-CoV-2 infection via various routes. K18-ACE2 mice were inoculated with approximately 10^4^ PFU SARS-CoV-2 via five different injection routes (n = 8 per injection route). Body weight (Left) and survival (Right) were monitored at the indicated dpi. SARS-CoV-2: severe acute respiratory syndrome coronavirus 2; PFU: plaque-forming unit; dpi: days post-infection

## Notes

### Competing Interest Statement

The authors have declared no competing interest.

## References

Azzolini, C., S. Donati, E. Premi, A. Baj, C. Siracusa, A. Genoni, P.A. Grossi, L. Azzi, F. Sessa, and F. Dentali. 2021. SARS-CoV-2 on ocular surfaces in a cohort of patients with COVID-19 from the lombardy region, Italy. JAMA ophthalmology 139:956–963.

Belser, J.A., P.A. Rota, and T.M. Tumpey. 2013. Ocular tropism of respiratory viruses. Microbiology and Molecular Biology Reviews 77:144–156.

Bojsen-Møller, F. 1975. Demonstration of terminalis, olfactory, trigeminal and perivascular nerves in the rat nasal septum. Journal of Comparative Neurology 159:245–256.

Chan, J.F.-W., A.J. Zhang, S. Yuan, V.K.-M. Poon, C.C.-S. Chan, A.C.-Y. Lee, W.-M. Chan, Z. Fan, H.-W. Tsoi, and L. Wen. 2020. Simulation of the clinical and pathological manifestations of coronavirus disease 2019 (COVID-19) in a golden Syrian hamster model: implications for disease pathogenesis and transmissibility. Clinical infectious diseases 71:2428–2446.

Chen, L., C. Deng, X. Chen, X. Zhang, B. Chen, H. Yu, Y. Qin, K. Xiao, H. Zhang, and X. Sun. 2020. Ocular manifestations and clinical characteristics of 535 cases of COVID-19 in Wuhan, China: a cross-sectional study. Acta ophthalmologica 98:e951–e959.

Colavita, F., D. Lapa, F. Carletti, E. Lalle, L. Bordi, P. Marsella, E. Nicastri, N. Bevilacqua, M.L. Giancola, and A. Corpolongo. 2020. SARS-CoV-2 isolation from ocular secretions of a patient with COVID-19 in Italy with prolonged viral RNA detection. Annals of internal medicine 173:242–243.

Davis, B.D., Dulbecco R., Eisen, H.N., Ginsberg, H.S., and Wood, W.B. 1972. “Nature of viruses” in Microbiology. (New York: Harper and Row) p. 1044–1053.

de Freitas Santoro, D., L.B. De Sousa, N.O. Câmara, D. De Freitas, and L.A. De Oliveira. 2021. SARS-COV-2 and ocular surface: from physiology to pathology, a route to understand transmission and disease. Frontiers in Physiology 12:106.

de Melo, G.D., F. Lazarini, S. Levallois, C. Hautefort, V. Michel, F. Larrous, B. Verillaud, C. Aparicio, S. Wagner, and G. Gheusi. 2021. COVID-19–related anosmia is associated with viral persistence and inflammation in human olfactory epithelium and brain infection in hamsters. Science Translational Medicine 13:eabf8396.

Deng, W., L. Bao, H. Gao, Z. Xiang, Y. Qu, Z. Song, S. Gong, J. Liu, J. Liu, and P. Yu. 2020. Ocular conjunctival inoculation of SARS-CoV-2 can cause mild COVID-19 in rhesus macaques. Nature communications 11:1–7.

DuPage, M., A.L. Dooley, and T. Jacks. 2009. Conditional mouse lung cancer models using adenoviral or lentiviral delivery of Cre recombinase. Nature Protocols 4:1064–1072.

Eriksen, A.Z., R. Møller, B. Makovoz, S.A. Uhl, and T.A. Blenkinsop. 2021. SARS-CoV-2 infects human adult donor eyes and hESC-derived ocular epithelium. Cell stem cell 28:1205–1220. e1207.

Fox, M. 1965. The visual cliff test for the study of visual depth perception in the mouse. Animal behaviour 13:232–IN233.

Gasparini, M.S., L.M. Dos Santos, A.M. Hamade, L.G. Gross, A.P. Favarato, J.P. de Vasconcellos, M.B. de Melo, P.L. Parise, C.L. Simeoni, and N.B. Silva. 2021. Identification of SARS-CoV-2 on the ocular surface in a cohort of COVID-19 patients from Brazil. Experimental Biology and Medicine 246:2495–2501.

Hassan, A.O., J.B. Case, E.S. Winkler, L.B. Thackray, N.M. Kafai, A.L. Bailey, B.T. McCune, J.M. Fox, R.E. Chen, and W.B. Alsoussi. 2020. A SARS-CoV-2 infection model in mice demonstrates protection by neutralizing antibodies. Cell 182:744–753. e744.

Jureka A.S., Silvas J.A and Basler C.F. 2020. Propagation, Inactivation, and Safety Testing of SARS-CoV-2. Viruses 12(6):622.

Imai, M., K. Iwatsuki-Horimoto, M. Hatta, S. Loeber, P.J. Halfmann, N. Nakajima, T. Watanabe, M. Ujie, K. Takahashi, and M. Ito. 2020. Syrian hamsters as a small animal model for SARS-CoV-2 infection and countermeasure development. Proceedings of the National Academy of Sciences 117:16587–16595.

Karimi, S., A. Arabi, T. Shahraki, and S. Safi. 2020. Detection of severe acute respiratory syndrome Coronavirus-2 in the tears of patients with Coronavirus disease 2019. Eye 34:1220–1223.

Leonardi, A., U. Rosani, and P. Brun. 2020. Ocular surface expression of SARS-CoV-2 receptors. Ocular immunology and inflammation 28:735–738.

Li, M., Y. Yang, T. He, R. Wei, Y. Shen, T. Qi, T. Han, Z. Song, Z. Zhu, and X. Ma. 2021. Detection of SARS-CoV-2 in the ocular surface in different phases of COVID-19 patients in Shanghai, China. Annals of Translational Medicine 9:

List, W., P. Regitnig, K. Kashofer, G. Gorkiewicz, M. Zacharias, A. Wedrich, and L. Posch-Pertl. 2020. Occurrence of SARS-CoV-2 in the intraocular milieu. Experimental Eye Research 201:108273.

Liu, T., J. Li, L. Yu, H.-X. Sun, J. Li, G. Dong, Y. Hu, Y. Li, Y. Shen, and J. Wu. 2021a. Cross-species single-cell transcriptomic analysis reveals pre-gastrulation developmental differences among pigs, monkeys, and humans. Cell discovery 7:1–17.

Liu, X., A. Zaid, J.R. Freitas, N.A. McMillan, S. Mahalingam, and A. Taylor. 2021b. Infectious clones produce SARS-CoV-2 that causes severe pulmonary disease in infected K18-human ACE2 mice. Mbio 12:e00819–00821.

Lochhead, J.J., and T.P. Davis. 2019. Perivascular and perineural pathways involved in brain delivery and distribution of drugs after intranasal administration. Pharmaceutics 11:598.

Loon, S., S. Teoh, L. Oon, S. Se-Thoe, A. Ling, Y. Leo, and H. Leong. 2004. The severe acute respiratory syndrome coronavirus in tears. British journal of ophthalmology 88:861–863.

Luan, J., Y. Lu, X. Jin, and L. Zhang. 2020. Spike protein recognition of mammalian ACE2 predicts the host range and an optimized ACE2 for SARS-CoV-2 infection. Biochemical and biophysical research communications 526:165–169.

McCray Jr, P.B., L. Pewe, C. Wohlford-Lenane, M. Hickey, L. Manzel, L. Shi, J. Netland, H.P. Jia, C. Halabi, and C.D. Sigmund. 2007. Lethal infection of K18-hACE2 mice infected with severe acute respiratory syndrome coronavirus. Journal of virology 81:813–821.

Meinhardt, J., J. Radke, C. Dittmayer, J. Franz, C. Thomas, R. Mothes, M. Laue, J. Schneider, S. Brünink, and S. Greuel. 2021. Olfactory transmucosal SARS-CoV-2 invasion as a port of central nervous system entry in individuals with COVID-19. Nature neuroscience 24:168–175.

Miner, J.J., A. Sene, J.M. Richner, A.M. Smith, A. Santeford, N. Ban, J. Weger-Lucarelli, F. Manzella, C. Rückert, and J. Govero. 2016. Zika virus infection in mice causes panuveitis with shedding of virus in tears. Cell reports 16:3208–3218.

Molina-Gil, J., L. González-Fernández, and C. García-Cabo. 2021. Trigeminal neuralgia as the sole neurological manifestation of COVID-19: A case report. Headache: The Journal of Head and Face Pain 61:560–562.

Netland, J., D.K. Meyerholz, S. Moore, M. Cassell, and S. Perlman. 2008. Severe acute respiratory syndrome coronavirus infection causes neuronal death in the absence of encephalitis in mice transgenic for human ACE2. Journal of virology 82:7264–7275.

Norwood, J.N., Q. Zhang, D. Card, A. Craine, T.M. Ryan, and P.J. Drew. 2019. Anatomical basis and physiological role of cerebrospinal fluid transport through the murine cribriform plate. Elife 8:e44278.

Penkava, J., M. Muenchhoff, I. Badell, A. Osterman, C. Delbridge, F. Niederbuchner, S. Soliman, M. Rudelius, A. Graf, and S. Krebs. 2021. Detection of SARS-CoV-2-RNA in post-mortem samples of human eyes. Graefe’s Archive for Clinical and Experimental Ophthalmology 1–9.

Pirraglia, M.P., G. Ceccarelli, A. Cerini, G. Visioli, G. d’Ettorre, C.M. Mastroianni, F. Pugliese, A. Lambiase, and M. Gharbiya. 2020. Retinal involvement and ocular findings in COVID-19 pneumonia patients. Scientific Reports 10:1–7.

Puelles, V.G., M. Lütgehetmann, M.T. Lindenmeyer, J.P. Sperhake, M.N. Wong, L. Allweiss, S. Chilla, A. Heinemann, N. Wanner, and S. Liu. 2020. Multiorgan and renal tropism of SARS-CoV-2. New England Journal of Medicine 383:590–592.

Ramani, A., A.-I. Pranty, and J. Gopalakrishnan. 2021. Neurotropic effects of SARS-CoV-2 modeled by the human brain organoids. Stem Cell Reports 16:373–384.

Reed, L.J. and Muench, H. 1938. A simple method of estimating fifty per cent endpoints. Am. J. Epidemiol. 27:493–497.

Rihn, S.J., A. Merits, S. Bakshi, M.L. Turnbull, A. Wickenhagen, A.J. Alexander, C. Baillie, B. Brennan, F. Brown, and K. Brunker. 2021. A plasmid DNA-launched SARS-CoV-2 reverse genetics system and coronavirus toolkit for COVID-19 research. PLoS biology 19:e3001091.

Sawant, O.B., S. Singh, R.E. Wright III, K.M. Jones, M.S. Titus, E. Dennis, E. Hicks, P.A. Majmudar, A. Kumar, and S.I. Mian. 2021. Prevalence of SARS-CoV-2 in human post-mortem ocular tissues. The Ocular Surface 19:322–329.

Schaefer, M.L., B. Böttger, W.L. Silver, and T.E. Finger. 2002. Trigeminal collaterals in the nasal epithelium and olfactory bulb: a potential route for direct modulation of olfactory information by trigeminal stimuli. Journal of Comparative Neurology 444:221–226.

Shimizu, S. 2004. Routes of administration. The laboratory mouse 1:527–543.

Sia, S.F., L.-M. Yan, A.W. Chin, K. Fung, K.-T. Choy, A.Y. Wong, P. Kaewpreedee, R.A. Perera, L.L. Poon, and J.M. Nicholls. 2020. Pathogenesis and transmission of SARS-CoV-2 in golden hamsters. Nature 583:834–838.

Singh, S., G. Garcia Jr, R. Shah, A.A. Kramerov, R.E. Wright III, T.M. Spektor, A.V. Ljubimov, V. Arumugaswami, and A. Kumar. 2021. SARS-CoV-2 and its beta variant of concern infect human conjunctival epithelial cells and induce differential antiviral innate immune response. The ocular surface

Song, E., C. Zhang, B. Israelow, A. Lu-Culligan, A.V. Prado, S. Skriabine, P. Lu, O.-E. Weizman, F. Liu, and Y. Dai. 2021. Neuroinvasion of SARS-CoV-2 in human and mouse brain. Journal of Experimental Medicine 218:

Steiner, M., M. del Mar Esteban-Ortega, and S. Muñoz-Fernández. 2019. Choroidal and retinal thickness in systemic autoimmune and inflammatory diseases: a review. Survey of ophthalmology 64:757–769.

Thakur, V., R.K. Ratho, P. Kumar, S.K. Bhatia, I. Bora, G.K. Mohi, S.K. Saxena, M. Devi, D. Yadav, and S. Mehariya. 2021. Multi-organ involvement in COVID-19: beyond pulmonary manifestations. Journal of clinical medicine 10:446.

Tzameret, A., I. Sher, M. Belkin, A.J. Treves, A. Meir, A. Nagler, H. Levkovitch-Verbin, Y. Rotenstreich, and A.S. Solomon. 2015. Epiretinal transplantation of human bone marrow mesenchymal stem cells rescues retinal and vision function in a rat model of retinal degeneration. Stem Cell Research 15:387–394.

Wan, K.H., S.S. Huang, and D.S. Lam. 2021. Conjunctival findings in patients with coronavirus disease 2019. JAMA ophthalmology 139:254–255.

Willcox, M.D., K. Walsh, J.J. Nichols, P.B. Morgan, and L.W. Jones. 2020. The ocular surface, coronaviruses and COVID-19. Clinical and Experimental Optometry 103:418–424.

Winkler, E.S., A.L. Bailey, N.M. Kafai, S. Nair, B.T. McCune, J. Yu, J.M. Fox, R.E. Chen, J.T. Earnest, and S.P. Keeler. 2020. SARS-CoV-2 infection of human ACE2-transgenic mice causes severe lung inflammation and impaired function. Nature immunology 21:1327–1335.

Yan, Y., B. Diao, Y. Liu, W. Zhang, G. Wang, and X. Chen. 2020. Severe acute respiratory syndrome coronavirus 2 nucleocapsid protein in the ocular tissues of a patient previously infected with coronavirus disease 2019. JAMA ophthalmology 138:1201–1204.

Zhang, A.J., A.C.-Y. Lee, H. Chu, J.F.-W. Chan, Z. Fan, C. Li, F. Liu, Y. Chen, S. Yuan, and V.K.-M. Poon. 2021. Severe acute respiratory syndrome coronavirus 2 infects and damages the mature and immature olfactory sensory neurons of hamsters. Clinical infectious diseases 73:e503–e512.

Zhou, L., Z. Xu, G.M. Castiglione, U.S. Soiberman, C.G. Eberhart, and E.J. Duh. 2020a. ACE2 and TMPRSS2 are expressed on the human ocular surface, suggesting susceptibility to SARS-CoV-2 infection. The ocular surface 18:537–544.

Zhou, Z., H. Kang, S. Li, and X. Zhao. 2020b. Understanding the neurotropic characteristics of SARS-CoV-2: from neurological manifestations of COVID-19 to potential neurotropic mechanisms. Journal of neurology 267:2179–2184.

